# Carnelian: alignment-free functional binning and abundance estimation of metagenomic reads

**DOI:** 10.1101/375121

**Authors:** Sumaiya Nazeen, Bonnie Berger

## Abstract

Accurate assignment of metagenomic reads to their functional roles is an important first step towards gaining insights into the relationship between the human microbiomeincluding the collective genesand disease. Existing approaches focus on binning sequencing reads into known taxonomic classes or by genes, often failing to produce results that generalize across different cohorts with the same disease. We present *Carnelian*, a highly precise and accurate pipeline for alignment-free functional binning and abundance estimation, which leverages the recent idea of *even-coverage*, *low-density* locality sensitive hashing. When coupled with one-against-all classifiers, reads can be binned by molecular function encoded in their gene content with higher precision and accuracy. *Carnelians minutes-per-metagenome* processing speed enables analysis of large-scale disease or environmental datasets to reveal disease- and environment-specific changes in microbial functionality previously poorly understood. Our pipeline newly reveals a functional dysbiosis in patient gut microbiomes, not found in earlier metagenomic studies, and identifies a distinct shift from matched healthy individuals in Type-2 Diabetes (T2D) and early-stage Parkinson’s Disease (PD). We remarkably identify a set of functional markers that can differentiate between patients and healthy individuals consistently across both the datasets with high specificity.

## 1 Introduction

Metagenomics fundamentally asks what microorganisms are present in a sample, what functions they perform, and how they compare across different conditions. However, the sequencing datasets required to answer these questions are gigantic and incredibly complex since they come from a mixture of many different, sometimes related organisms as opposed to standard genomic datasets which come from a single organism. This data results in major identification challenges which are complicated further by the fact that the data is from diverse origins, has low coverage, contains a high rate of sequencing errors, and typically has short sequencing reads. Taken together, these challenges drive the need for clever algorithms and efficient tools in order to draw meaningful insights from the data. While classical metagenomic binning asks what organisms are present in a sample (taxonomic classification), functional binning focuses on annotating metagenomic reads with functional labels (usually from gene content) and estimating their relative abundance in the dataset. This in turn provides information about the abundance of biochemical properties and metabolic pathways in the dataset. Such knowledge is imperative to our overall understanding of the precise functional relationships between the human gut microbiome and different diseases. Over the years, hundreds of clinical studies have demonstrated associations between the human microbiome and disease characterized by pathogenic dysbiosis in patient microbiomes (1). Some diseases have been marked by an invasion of harmful pathogens, whereas others have been marked by a depletion of beneficial ones (2). For example, the human gut microbiome has been found to be associated with Crohn’s Disease (3), obesity (4), Type-2 Diabetes (5; 6), Colorectal Cancer (7), Parkinson’s Disease (8; 9) and even Autism Spectrum Disorder which is found to have an innate immunity component (10; 11; 12). However, the precise workings of human microbiome in different diseases remain poorly understood till date.

Classical approaches for functional binning perform genome assembly where the reads are assembled into large contigs and annotated using sequence homology, often using existing alignment tools such as BLAST (13), profile Hidden Markov Models (HMMs) or position-specific weight matrices (PWMs). Examples of such methods include, RAST (14), Megan4 (15), MEDUSA (16), Tentacle (17), MOCat2 (18), IMG4 (19), and so on. However, assembly is a slow, resource-heavy and lossy process (even with a fast aligner such as CORA (20)), especially for complex and diverse metagenomic samples. The lack of scalability and precision makes downstream gene finding and abundance estimation difficult on such data. Several platforms perform functional binning of reads directly via sequence homology, such as MG-RAST (21), but suffer from low precision due to the short read length and lack of high-quality gold standard references. Also, alignment of reads is a resource-heavy and time consuming process which becomes prohibitive while comparing hundreds of samples from different disease conditions (22; 23). Recently published *mi-faser* (24) speeds up the alignment step significantly by using a fast and precise protein aligner, DIAMOND (25) in a multi-node cluster environment. All alignment-based methods annotate only a fraction of the reads and suffer from low sensitivity.

On the other end, there are the faster compositional approaches which perform alignment-free taxonomic binning of reads according to specific patterns of their constituent k-mers (e.g., Kraken (26) and CLARK (22). They are very efficient but limited in terms of sensitivity. Metakallisto (27), a pseudo-alignment technique for metagenomic taxonomic classification, relies on the idea of approximate k-mer matching via a k-mer de Bruijn graph – an idea originally introduced by Kallisto (28) for quantifying transcript abundances in RNAseq data. Other approaches rely on supervised machine learning (ML) classifiers, such as, Naïve Bayes (NB) or support vector machines (SVM), trained on a set of reference sequences to classify taxonomic origins of metagenomic reads using their relative k-mer frequency vectors (29; 30). These approaches are much faster than alignment-based methods but need a *high-density* k-mer representation thus a much larger feature space. Opal (31), a tool for metagenomic taxonomic binning, reduces the size of feature space by orders of magnitude via *low-density* even-coverage locality sensitive hashing (LSH), inspired by Gallager’s low density parity check (LDPC) codes (32). We build upon this idea of Opal-Gallager hashes to perform *functional* metagenomic binning (versus Opal’s taxonomic binning) more accurately in the presence of only a moderate degree of sequence homology.

We present *Carnelian*, an alignment-free functional binning and abundance estimation pipeline, which uses Opal-Gallager hashes (31) to represent translated metagenomic reads (amino-acid sequences) into low-dimensional manifolds to construct a compact feature space, then functionally bin them with a set of one-against-all classifiers. The classifier ensemble is trained on quality annotated gene sequences from a novel curated gold standard reference with experimentally validated functional labels. *Carnelian* achieves higher accuracy (F1-score) in binning both “seen” (98.26%) and “unseen” proteins (72.53%) into functional bins (Enzyme-commission labels) as compared to state-of-the art methods: mi-faser and Kallisto. On a single CPU core, *Carnelian* requires ∼ 11 minutes to annotate ∼ 1.5 million translated contigs of ∼ 100 amino acids (AA) each and ∼ 59 minutes to annotate ∼ 8.6 million translated read fragments of ∼ 30 AA each from a human fecal sample.

When applied to two large Type-2 Diabetes (T2D) case-control studies of Chinese (5) and European (16) population, our pipeline newly reveals a functional dysbiosis in the T2D patient gut; in particular, *Carnelian* differentiates T2D patients from healthy controls in terms of 14 shared dysregulated metabolic pathways, many of which are not found using mi-faser. Remarkably, we identify a novel set of functional markers that can classify T2D patients versus healthy individuals consistently across datasets with higher specificity (AUC ∼ 78%). When run on a Parkinsons Disease (PD) dataset, *Carnelian* highlights the altered microbial functionality in early-stage PD patient guts in terms of increased lipid metabolism, and reduced glucose metabolism, and tryptophan biosynthesis — none of these found by the original case-control study (33). Notably, *Carnelian*-generated functional profiles capture more complete information than mi-faser and enable us to reveal biologically relevant pathways not found using mi-faser profiles.

## 2 Results and Discussion

### *Carnelian* pipeline

Current alignment-based and compositional methods focus on achieving only high precision and suffer from low sensitivity. We introduce a highly precise and sensitive pipeline Carnelian that: (i) translates reads (amino acid sequences) from whole metagenome sequencing studies; (ii) leverages the low-density even-coverage Opal-Gallager hashes (31; 32) to encode translated reads into low-dimensional manifolds to produce a compact feature vector representation; and (iii) employs an ensemble of one-against-all classifiers to bin the compact feature vectors by metabolic function (versus taxa as with OPAL) to produce functional vectors containing effective read counts i.e. read counts normalized against average gene length per functional label, average read length, as well as total number of annotated fragments per label. The intuition behind this is similar to transcripts per million (TPM) counts used for RNAseq data (34). Effective read counts can be fed into any standard abundance estimation pipeline designed for count data to produce functional abundance estimations. Advantageously, our compact feature vectors make Carnelian robust against sequencing errors and reduce the size of the feature space compared to other compositional methods, which saves memory. We chose to train Vowpal-Wabbit one-against-all classifiers (35) with the compact feature vectors of protein sequences from our in-house curated gold standard dataset in an online fashion, which requires only one training sample in memory at a time; such classifiers make Carnelian incrementally trainable and memory efficient. Figure 1 depicts the main steps of the *Carnelian* pipeline.

**Figure 1:**
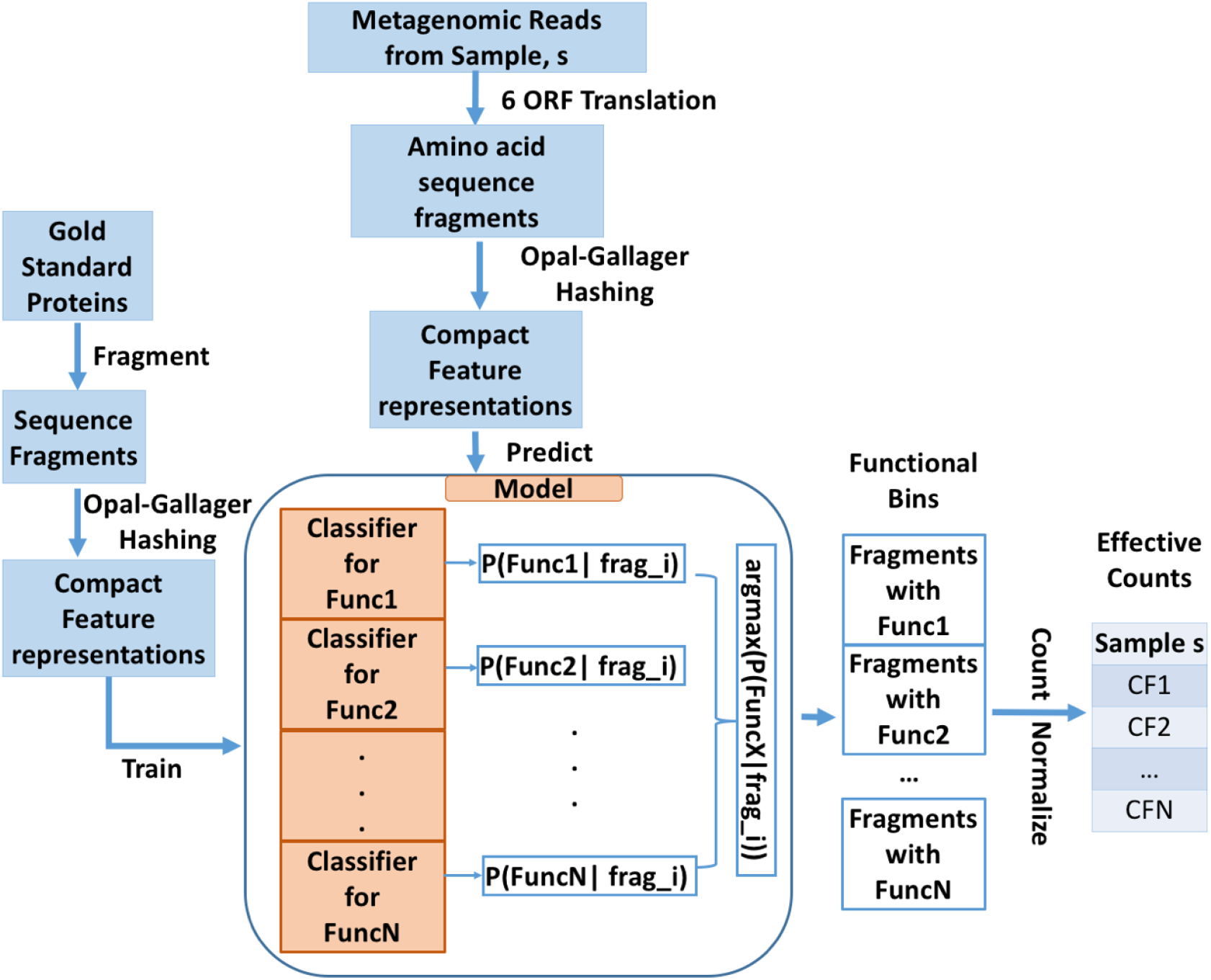
Carnelian pipeline. *Carnelian* trains on functionally annotated protein sequences by generating fixed-length (*l*) fragments and their low-density spaced k-mer representations which are used as features by a set of one-against-all online classifiers. The learned model is then used to bin input amino acid sequences into appropriate functional bins. Abundance estimates are constructed from effective fragment counts in functional bins for each sample, and downstream differential abundance analysis is performed to find dysregulated ECs and pathways.

Since cheaper Amplicon-based sequencing techniques focus on the diversity of specific marker genes (e.g., the 16S rRNA gene) and are limited in their ability to infer molecular function, we designed *Carnelian* for analyzing whole metagenome sequencing data (e.g., Illumina HiSeq (36)). Unfortunately, this data comes from a mixture of many different and sometimes related organisms and can encode a 100-times more unique genes than ones own genome (37; 38). Moreover, only a small fraction of these genes have known functional annotations. Thus, in our choice of gold standard functional annotations, we restricted ourselves to proteins that have unique and complete Enzyme Commission (EC) numbers associated with them.

### *Carnelian* outperforms state-of-the-art methods

*Carnelian* achieves high accuracy in binning both “seen” proteins and “unseen” proteins into EC labels, a numerical classification scheme for enzymes, based on the chemical reactions they catalyze (see Methods). On our in-house gold standard dataset, EC-2192-DB, *Carnelian* achieves almost an F1-score of 98.26% while annotating randomly drawn fragments from “seen” proteins (Table 1). Moreover, it achieves an F1-score of 72.53% and demonstrates greater sensitivity to novel proteins while annotating fragments from proteins not seen by the classifiers during training (Figure 2). In both cases, *Carnelian* outperforms Kallisto and mi-faser in terms of F1-score. Both Kallisto and mi-faser do not annotate a significant portion of the test fragments in case of “unseen” proteins, which indicates the richness of the spaced k-mer representation learned by *Carnelian*. *Carnelian* in its “default” mode emphasizes achieving both precision and recall per functional bin and enables the user to opt for “precise” mode when high-precision annotation is required; these features make it a more flexible tool of choice. Although, *Carnelian* uses more memory than mi-faser and Kallisto for the EC-2192-DB dataset, the memory requirement can be easily reduced by using a reduced amino acid alphabet (Supplementary Table T1). On smaller training sets Kallisto is the fastest and most memory efficient tool, however, as the training set becomes larger, its memory requirement exponentially increases and speed decreases substantially, while Carnelian still remains reasonably speedy and memory efficient (Supplementary Table T2).

**Table 1:** Performance comparison between Carnelian, mi-faser, and Kallisto

| Method | Sensitivity<br>(%) | Precision<br>(%) | F1-score<br>(%) | Speed<br>(frags / sec) | Peak Memory<br>(GB) |
| --- | --- | --- | --- | --- | --- |
| Carnelian | **98.86** | 97.86 | **98.26** | 2446.76 | 6.43 |
| Carnelian (precise) | 76.10 | 99.80 | 85.52 | 2313.10 | 6.43 |
| mi-faser | 96.78 | **99.95** | 98.16 | 1880.37 | 1.31 |
| Kallisto | 90.33 | 99.67 | 93.90 | **2786.75** | **0.42** |

**Figure 2:**
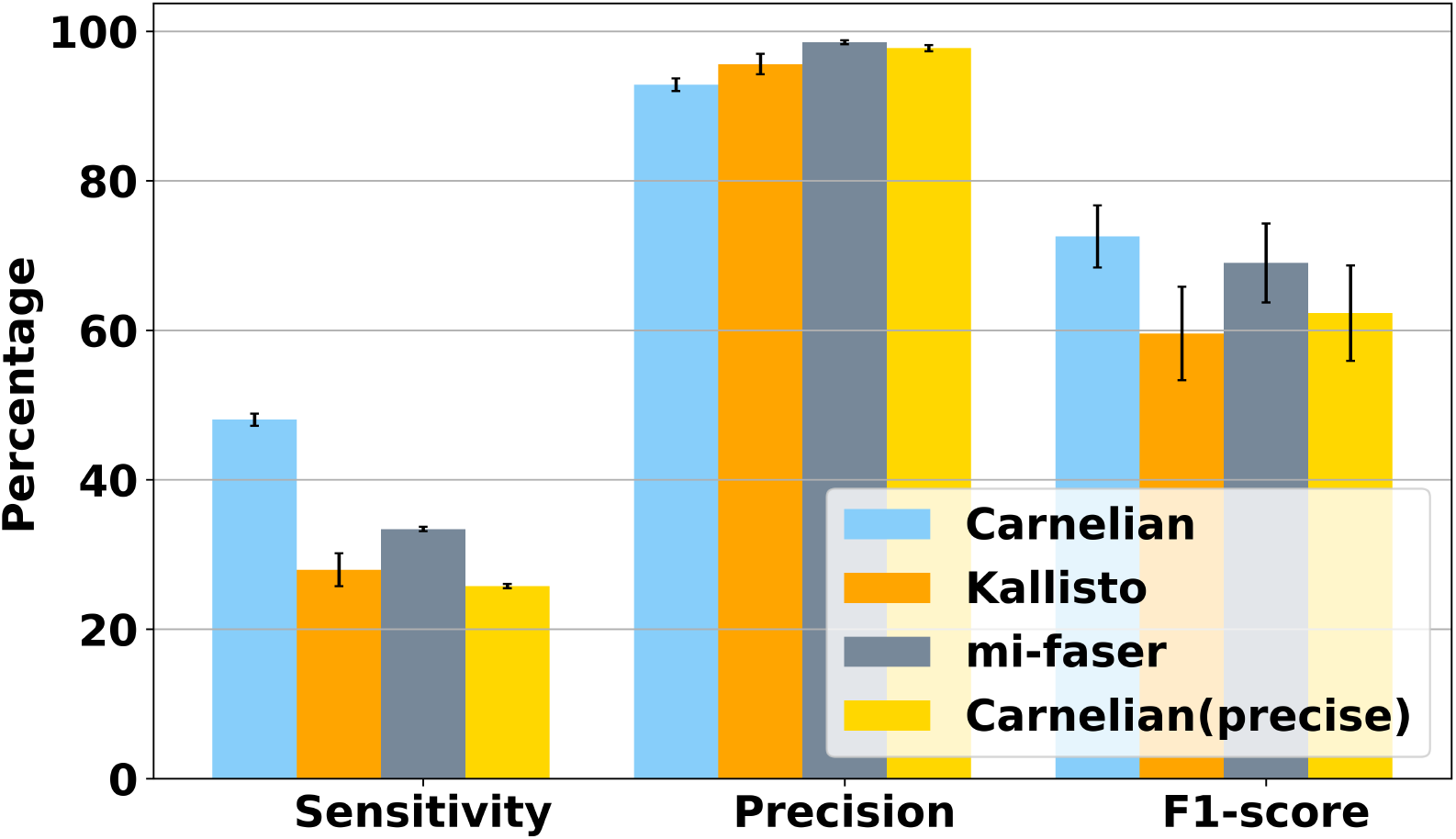
Binning performance of *Carnelian* on novel proteins. In “default” mode, *Carnelian* achieves much better sensitivity and F1-score than both Kallisto and mi-faser. In “precise” mode with a probability cut-off of 80%, *Carnelian* achieves comparable precision as Kallisto and mi-faser.

### *Carnelian* reveals T2D-related functional dysbiosis in gut microbiota

Using *Carnelian* pipeline we analyzed whole metagenome sequencing reads from the fecal samples of 145 Chinese individuals in the T2D-Qin dataset (5) and revealed microbial functional dysbiosis in T2D patients versus healthy controls. All healthy individuals in T2D-Qin dataset shared highly similar functional profiles (*r* = 0.99 ± 0.01) whereas functional profiles of patients with known disease status were more variable (*r* = 0.73±0.18, *p* = 0.0023). We identified 33 differentially altered enzymes between female T2D patients and controls and 7 such enzymes between male patients and controls using a log-fold-change threshold either less than −.58 (fold-change 0.67) or greater than 0.58 (fold change 1.5) and FDR corrected *p*-value cutoff 0.1 (Figure 3(a) & 3(b) and Supplementary Table T3). We mapped these enzymes to 46 metabolic pathways in the KEGG database (Supplementary Table T4). Among these pathways four were altered in the guts of both T2D males and females, namely glutathione metabolism, lysine biosynthesis, butanoate/butyrate metabolism, and tryosine metabolism. Dysregulation of these pathways have been implicated in T2D pathogenesis in earlier clinical studies (39; 40; 41; 42) and have therapeutic potential to improve insulin sensitivity in T2D patients.

**Figure 3:**
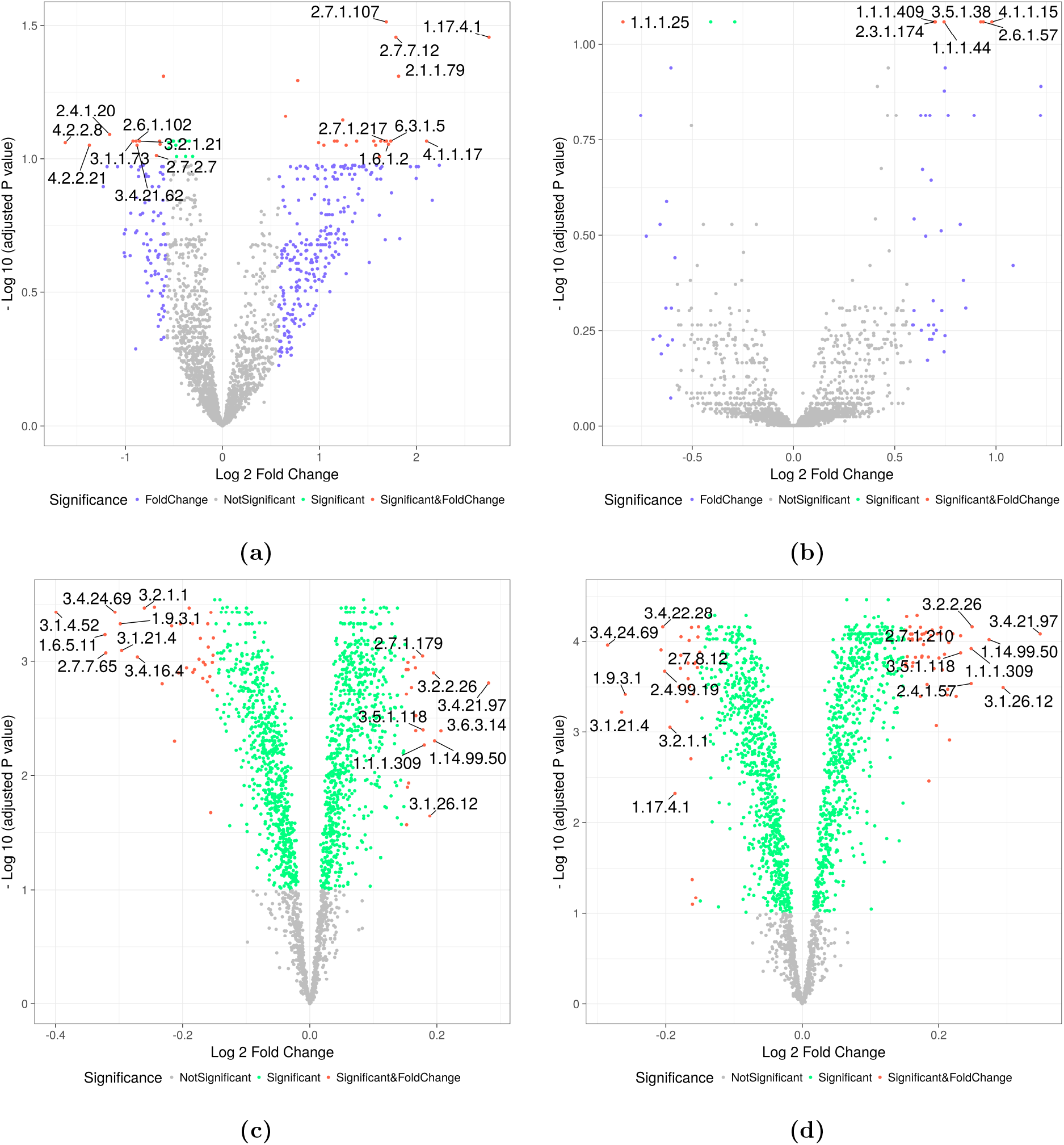
Differentially abundant/depleted ECs in T2D studies identified by *Carnelian*. Female T2D patients versus female controls in T2D-Qin dataset; **(b)** Male T2D patients versus male controls in T2D-Qin dataset. **(c)** T2D patients versus NGT individuals in T2D-Karlsson dataset; **(d)** T2D patients versus IGT individuals in T2D-Karlsson dataset.

In particular, we found Chinese patient guts to be enriched in a number of oxidoreductases and NAD+ synthase which may indicate higher degree of bacterial defense against oxidative stress — this finding is in agreement with the original study (5). There was an enrichment of transferases involved in amino acid and short chain fatty acid synthesis and energy homeostasis. Greater energy imbalance and enrichment of amino acids and short chain fatty acids cause instability in a number of other metabolic pathways, including bile salt metabolism, steroid metabolism, and phospholipid metabolism, and have been implicated in the development of obesity, insulin resistance, and T2D (43). We also found reduced levels of multiple lyases, glycosyltransferases, transaminases, isomerases, and nucleotidyltransferases which are key players in starch, sugar, and amino sugar metabolism, fatty acid oxidation, and metabolism of vitamin B6, vitamin B12 and folates. Reduced absorption of folates and vitamin B could be a side-effect of prolonged use of metformin, which is the first choice drug in diabetes (44). The enrichment of LpxC enzyme which aids Lipopolysaccharide biosynthesis may contribute in developing insulin resistance (45). Furthermore, we saw an increase of ornithine decarboxylase which can increase the rate of glutathione metabolism in T2D patients. Reduced levels of glutathione has been observed in T2D patients in earlier clinical studies (40). Finally, an enrichment of enzymes that take part in drug metabolism is suggestive of the hostile nature of T2D patient guts.

To test the generalizability of *Carnelian* pipeline, we applied it on the European individuals in the T2D-Karlsson dataset. We identified 54 differentially abundant/depleted enzymes between T2D patients and the individuals with normal glucose tolerance (NGT) and 84 such enzymes between T2D patients and individuals with impaired glucose tolerance (IGT) using a log-fold-change threshold either less than −.15 (fold change 0.9) or greater than 0.15 (fold change 1.1) and FDR corrected *p*-value cutoff of 0.1 (Figure 3(c) & 3(d) and Supplementary Tables T5 and T6).

Similar to our finding in Chinese population, we saw an enrichment of oxidoreductases that are directly involved in oxidative phosphorylation, glycolysis, glutathione and energy metabolism, confirming that there is higher oxidative stress in the T2D gut. We also found altered levels of transferases involved in amino acid homeostasis, metabolism of lipids, glycans, vitamins and co-factors, and steroid hormone biosynthesis. Additionally, we found a number of hydrolases and lyases enriched in the patient gut that are key players in carbohydrate and energy metabolism. These findings are supported by the original study (6). Notably, we observed depletion of ribonucleoside-diphosphate reductase and enrichment of bacterial ribonuclease E. Ribonucleoside-diphosphate reductase converts ribonucleoside diphosphates into deoxyribonucleoside diphosphates — essential for DNA synthesis and repair; this may indicate oxidative stress related DNA damage in T2D patients. Enrichment of bacterial ribonuclease E which is commonly found in *Escherichia Coli* (E. coli) could be indicative of an enrichment of such opportunistic bacterial species in the T2D and IGT guts.

We mapped the significantly enriched/depleted enzymes in T2D vs. NGT and T2D vs. IGT contrasts to KEGG metabolic pathways and found 27 pathways altered in both contrasts (Supplementary Table T7). Notably, 14 of these pathways were also found in our analysis of the T2D-Qin dataset. The common pathways include: carbohydrate metabolism pathways such as starch and sucrose metabolism, glycolysis/gluconeogenesis, galactose metabolism, pyruvate metabolism, etc.; nucleotide metabolism pathways such as purine and pyrimidine metabolism; vitamin and co-factor metabolism pathways such as nicotinate and nicotinamide metabolism and novobiocin biosynthesis; glycerophospholipid metabolism; and drug metabolism pathways. These pathways have been implicated in T2D in multiple clinical studies as discussed before and have the potential of being actionable to improve insulin sensitivity in patients.

To compare our results with mi-faser’s results, we applied mi-faser on both T2D-Qin and T2D-Karlsson datasets to generate functional profiles of T2D patients and controls and performed differential analysis of enzyme abundances. There was little overlap between the enzymes identified by *Carnelian* and mi-faser, however at pathway level there was more overlap (See Supplementary Tables T8-T11 for details). For the T2D-Qin dataset, *Carnelian* and mi-faser capture the important pathways such as, carbohydrate metabolism, butanoate metabolism, purine and pyrimidine metabolism etc. However, for the T2D-Karlsson dataset, mi-faser only annotated a fraction of the reads and was unable to find any of the pathways related to carbohydrate metabolism and oxidative stress between T2D and NGT individuals. One reason for this might be that, when read fragments are long, as in case of T2D-Qin datset, both the methods can capture valuable information. But, when the reads are short, as in case of T2D-Karlsson dataset, *Carnelian*-generated functional profiles capture more functional information than mi-faser-generated ones, thus facilitate the identification of more relevant biological insights.

### *Carnelian* facilitates gut microbiota based T2D classification

To test the ability of gut microbial functional markers to discriminate T2D patients from controls in the T2D-Qin dataset, we used *Carnelian*-identified and mi-faser-identified enzyme markers as features and performed 10-fold cross validation on the individuals using random forest classifiers.

We were able to achieve an AUC of 0.81 (95% CI: 0.74 - 0.88) with *Carnelian*, whereas using mi-faser identified enzymes as features, we could achieve an AUC of only 0.69 (95% CI: 0.60 - 0.77). The higher AUC achieved using *Carnelian*-identified markers may indicate that *Carnelian* identifies enzymes/metabolites that are functionally more relevant to T2D pathogenesis. Figure 4(a) shows the ROC curves for both random forest models on T2D-Qin dataset. We repeated the same analysis on T2D-Karlsson dataset to classify T2D individuals against NGT individuals. We were able to achieve an AUC of 0.88 (95% CI: 0.80 - 0.96) with *Carnelian*; in contrast, using mi-faser identified ECs as features, we could achieve an AUC of only 0.82 (95% CI: 0.73 - 0.91). Figure 4(b) shows the ROC curves for both random forest models for the T2D-Karlsson dataset. To test for generalizability, we used enzymes identified by *Carnelian* from the T2D-Qin dataset to classify the functional profiles of T2D and NGT individuals from the T2D-Karlsson dataset and vice-versa, achieving an average AUC of 0.76 in the former case and an average AUC of 0.60 in the latter. Finally, we compiled a list of 80 enzyme markers from the 40 markers identified in the T2D-Qin dataset and the 42 markers identified in both T2D vs. NGT and T2D vs. IGT contrasts by *Carnelian* (Supplementary Table T12); using these as features we classified *Carnelian*-generated functional profiles of T2D patients and controls and consistently achieved ∼ 78% AUC across both datasets (Figure 4(c)).

**Figure 4:**
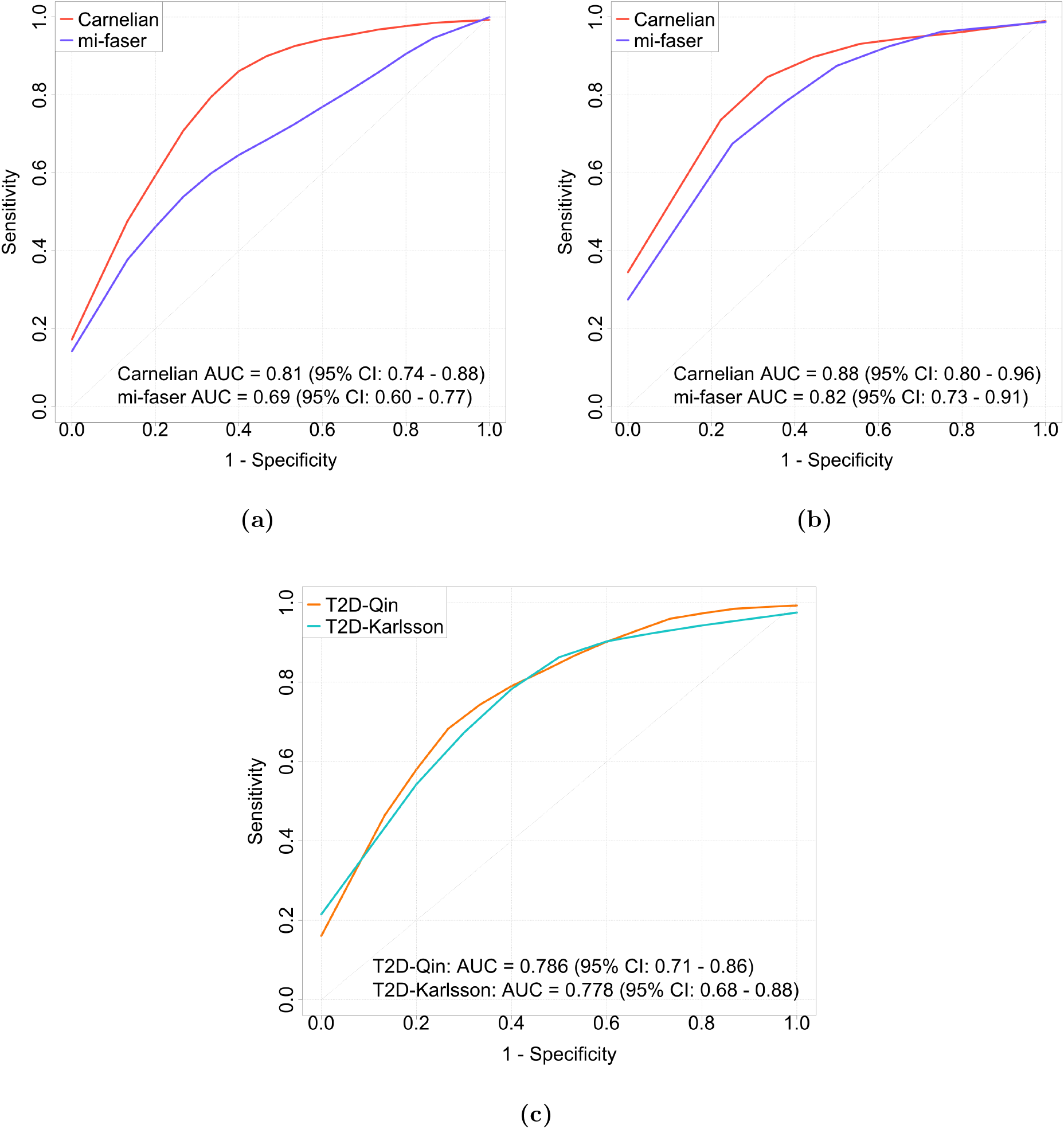
Calssification of T2D vs healthy using gut microbial functional markers. **(a)** T2D vs. normal in T2D-Qin dataset; **(b)** T2D vs. NGT in T2D-Karlsson dataset. **(c)** T2D vs. healthy using combined set of EC markers identified by *Carnelian*.

### *Carnelian* finds functional dysbiosis in PD patient gut microbiome

We applied *Carnelian* pipeline on whole metagenomic sequencing reads from the fecal samples of 20 early stage, L-DOPA naïve PD patients and 21 age-matched and gender-matched individuals in the PD-Bedarf dataset (33). Patients who had motor symptoms and diagnosis within the past year, and hadn’t been treated with L-DOPA, used to increase dopamine concentrations in the treatment of PD, were the target of the original study. Our analysis of this cohort revealed a distinct functional shift in the gut microbiome of early-stage PD patients from healthy controls. Using *Carnelian*-generated functional profiles, we identified 90 differentially enriched/depleted enzyme markers between PD patients and controls using a log-fold-change threshold either less than −.58 (fold change 0.67) or greater than 0.58 (fold change 1.5) and BH corrected *p*-value cutoff of 0.1 (Figure 5a and Supplementary Tables T13). These enzymes were mapped to 49 specific KEGG metabolic pathways (Supplementary Table T14).

**Figure 5:**
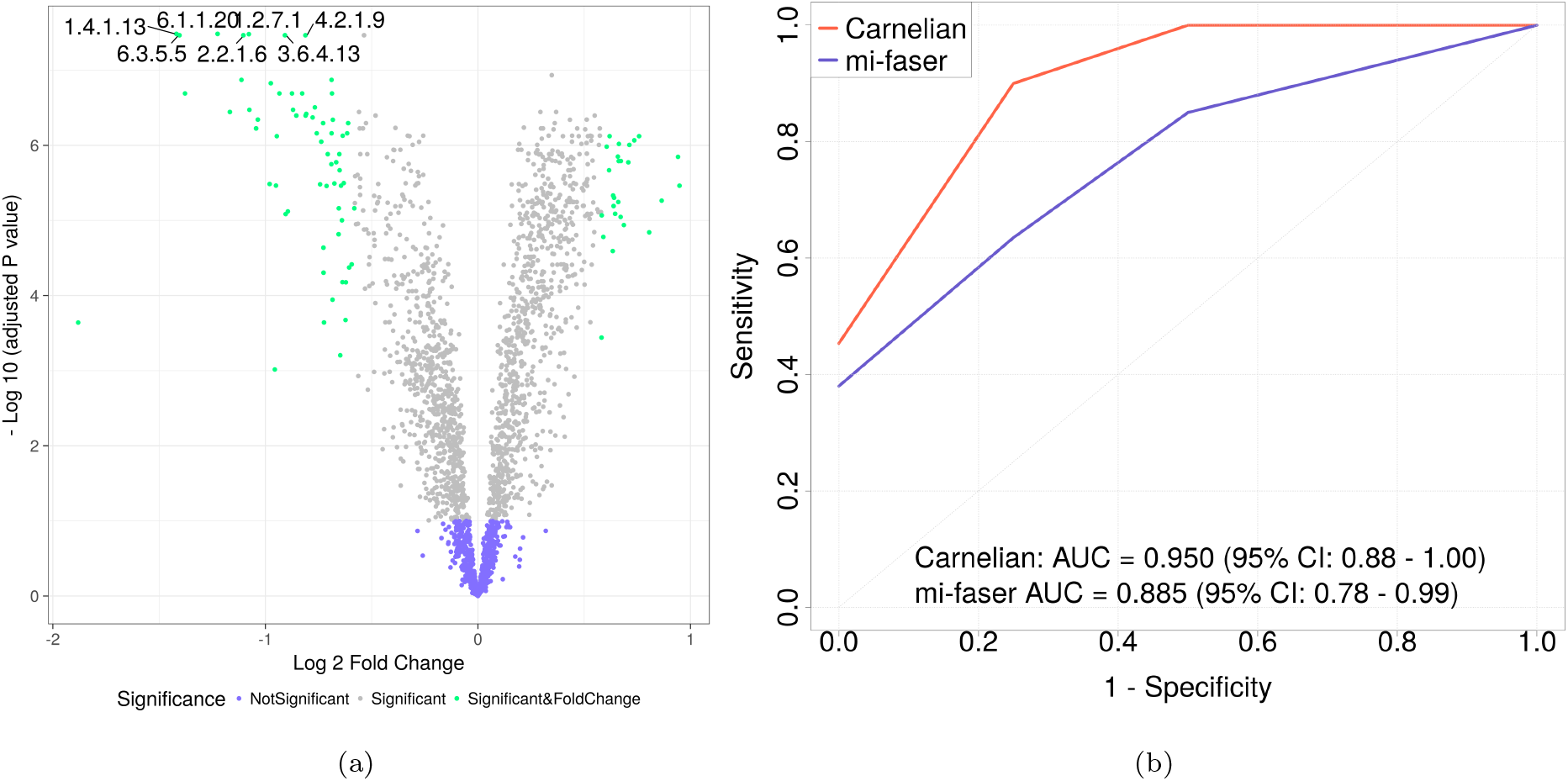
Analysis of PD-Bedarf dataset. **(a)** Differentially abundant/depleted EC labels identified by *Carnelian*. **(b)** Classification of PD vs healthy using EC markers identified by *Carnelian* and mi-faser.

We observed depletion of a large number of enzymes involved in carbohydrate metbolism (e.g., glycolysis/gluconeogenesis, starch and sucrose metabolism, starch and sucrose metabolism, pentose phosphate pathway). Disruption of glucose metabolism is known to be an early event in idiopathic PD (∼ 90% of all PD cases) and disruption of the pentose phosphate pathway is considered damaging to the neurons (46). We also observed altered levels (mainly depletion) of a number of oxidoreductases and transferases involved in energy metabolism which suggests a lower rate of energy metabolism and higher degree of oxidative stress in the PD patient guts. Oxidative stress is believed to be a common denominator for the dysfunction of a number of cellular processes that lead to the loss of dopaminergic neurons in PD (47). We also found a higher degree of glutathione metabolism which may contribute to lower levels of glutathione, a thiol tripeptide critical for protecting dopaminergic neurons. Restoring glutathione metabolism to normalcy may hold therapeutic benefits for PD patients (48).

Additionally, we found enrichment of a number of enzymes involved in lipid metabolism in the PD patient microbiomes. Dysregulation of lipid metabolism has been implicated in a number of neurodegenerative diseases including PD (49; 50). In particular, association of *α*-synuclein (a major constituent of Lewy bodies) with oxidized lipid metabolites can lead to mitochondrial dysfunction and ultimately loss of dopaminergic neurons (51). Finally, *Carnelian* generated functional profiles led us to a number of dysregulated amino acid metabolism pathways including phenylalanine, tyrosine and tryptophan biosynthesis, alanine, aspartate and glutamate metabolism etc., and purine (uric acid) metabolism pathway. Altered levels of alanine, tryptophan, and purine has been observed in earlier metabolite profiling studies of PD (52; 33; 53).

Repeating the same analysis with mi-faser generated functional profiles, we did not find any of the above pathways (Supplementary Tables T15 and T16). Notably, using the differentially abundant enzymes identified by *Carnelian* as features, we performed 5-fold cross validation using random forest classifiers and achieved an AUC of 0.95 (95% CI: 0.88 to 1.00) in discriminating between PD patients and healthy controls (Figure 5b).

## 3 Conclusion

In summary, we provide a standalone pipeline for alignment-free functional metagenomic binning and abundance estimation which we demonstrate facilitates highly accurate and precise binning of metagenomic reads across diverse microbial datasets. As experimentally-validated functional annotations for bacterial proteins become increasingly available, *Carnelian* can be incrementally trained with the newly available functions. Although *Carnelian* requires more memory than mi-faser as the size of the gold standard data grows, it can be easily remedied by using a reduced-size amino acid alphabet. When applied to metagenomic datasets from T2D and PD case-control cohorts, our pipeline provides a more complete picture of the functional dysbiosis in the patient microbiomes, which was previously poorly understood using taxonomic binning. While earlier studies showed only a moderate degree of taxonomic dysbiosis in the T2D gut that did not generalize across different datasets, we identify 14 common metabolic pathways — some with strong therapeutic potential — that are disrupted in both European and Chinese patient cohorts and explain the workings of those by enriched and depleted enzymatic actions. Moreover, *Carnelian* is able to find several clinically established hallmarks of PD (e.g., altered lipid and glucose homeostasis, lower levels of tryptophan, a precursor of serotonin) that were not found by earlier metagenomic studies of PD patients. *Carnelian* can construct more informative functional profiles by identifying functions of unseen proteins from rare bacterial species, and reveal interesting biological insights from the disease datasets. However, we suggest rigorous investigation into the implications of these findings, because, with the current smaller gold standard, *Carnelian* might overestimate read counts for well-studied protein functions. As more and more protein functions are experimentally verified, *Carnelian* will enable fast and more accurate binning of reads at higher functional levels, thus producing less noisy and more complete microbial functional profiles of microbial samples and facilitating downstream biological discovery. We expect *Carnelian* to be an essential component of the metagenomic analysis toolkit.

## 4 Materials and Methods

### Availability of *Carnelian* pipeline

Source code for *Carnelian* is available at http://carnelian.csail.mit.edu and https://github.com/snz20/carnelian.

### Datasets

We built our gold standard reference dataset by first collecting reviewed bacterial proteins from UniProtKB/Swiss-Prot (Feb. 2018) (54; 55) that have evidence of existence at either the protein or transcriptomic level and have explicit Enzyme Commission (EC) Numbers associated with them since EC numbers act as primary identifiers for metabolic pathway members. We excluded any protein that is inferred by homology or has incomplete EC as well as multiple EC annotations. We also collected catalytic residues with corresponding EC numbers and UniProt proteins for which a literature reference existed. We combined these two sets and removed redundant sequences, which gave us a reference dataset, EC-2192-DB, of 8,258 proteins with 2,192 unique EC numbers (available on the Carnelian website). Amino acid sequences for these proteins were downloaded from UniProt ((54).

To simulate the effect of the presence of a novel protein in a metagenome, we randomly removed a protein from each of 499 EC bins that contained more than five proteins from the EC-2192-DB dataset. This process left us with 7,759 protein sequences in the training set which we named EC-499-TR. Simulated fragments from the removed proteins were used as a test dataset, which was named as EC-499-TS.

Whole metagenome sequencing data on individual fecal samples from 71 cases (female = 25, male = 46) and 74 controls (female = 26, male = 48) were obtained from a study performed on Chinese T2D patients (5). We call this dataset T2D-Qin. Metadata for the samples is given in Supplementary Table T17. Raw paired-end Illumina reads are available publicly from NCBI short read archive (SRA) (Study accession: SRP008047). We downloaded high quality adapter free assembled contigs for all 145 samples, totaling *∼* 380 gigabases (Gb) from BGI’s repository (ftp://penguin.genomics.cn/pub/10.5524/100001_101000/100036/AssemblyContigs/–a large dataset.

Additionally, we analyzed fecal metagenome sequencing data from a T2D study performed on a European cohort of 145 women with either T2D or impaired glucose tolerance (IGT) or normal glucose tolerance (NGT) (6). We downloaded publicly available raw Illumina HiSeq 2000 paired-end reads from an NCBI SRA (Study accession: ERP002469); each individual metagenome contained ∼ 3 Gb on average. Metadata for the analyzed individuals is given in Supplementary Table T18. We used Trimmomatic v0.36 ((56) in paired-end mode for adapter trimming and quality filtering with a quality threshold of 30 and a minimum length of 90 bp. DeconSeq v0.4.3 (57) was used to remove contaminating human sequences with the human reference genome GRCh38 as database. We kept only the read-pairs for which both sequences survived quality control and named this dataset T2D-Karlsson. On average 75% of the read pairs were retained for downstream analysis.

We also analyzed whole metagenome sequencing reads from the fecal samples of 20 patients and 21 healthy individuals in a early stage L-DOPA naïve PD case-control study (33). All the participants in the study were male and age matched. Metadata of the participants in provided in Supplementary Table T19. Metagenomes were pre-Sprocessed in a similar fashion as T2D-Karlsson dataset. We called this dataset, PD-Bedarf.

### Constructing compact feature vectors using Opal-Gallager hashes

Let us consider a sequence fragment of *l* amino acids, *s* ∈ Σ^*l*^, where Σ = amino acid alphabet (*|*Σ*|* = 20). A k-mer, with *k < l*, is a short word of *k* contiguous amino acids. Similar to the bag-of-words representation of a document, we define a k-mer profile of a sequence *s* as a vector 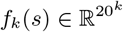. We index each k-mer with an integer *i*, where 0 *≤ i ≤* 20^*k*^ which can be represented by a binary string of length 5*k*. Each entry *f_k_*(*s, i*) ∈ *f_k_*(*s*) stores the frequency of the *i*-th k-mer. Thus, an amino acid fragment of length *l* can be represented using k-mers in *O*(20^*k*^) space instead of a vector of *O*(20^*l*^). Using random locality sensitive hash (LSH) functions, we can create k-mer profiles that specify spaced subsequences, rather than contiguous subsequences of fragment *s*. More specifically, we define a random hash function, *h* : Σ*^k^* → Σ*^r^* to generate a spaced (*k, r*)-mer such that a hashed k-mer can be represented by a binary vector of *O*(20^*r*^) dimensions with corresponding positions set to 1. With this family of LSH functions, we can randomly sample a set of *m* LSH functions and concatenate them together to represent a k-mer profile of a sequence by only *O*(*m*20^*r*^) *« O*(20^*k*^) space. However, k-mer profiles built with uniformly random LSH functions, often have low coverage of the beginning and end positions of a sequence in practice unless a large number of such functions are used. Our goal is to use a small number (*m*) of LSH functions that are sufficiently discriminating and informative for the *k*-mer in consideration which we achieve by building upon Opal’s modified Gallager design algorithm (31). Supplementary Figure F1 shows an example of how even coverage LSH functions are newly generated for an amino acid *k*-mer.

### Choice of fragment length and *k*-mer length

While choosing the fragment length, *l*, we needed to ensure that the fragments we draw are smaller than the smallest protein sequence in our datasets. Lengths of 8,258 protein sequences in EC-2192-DB range from 34 to 7,073 amino acids with a median length of 342 amino acids. Also, contigs from the T2D-Qin dataset have a minimum length of 166 amino acids, whereas the nominal length for paired end reads from the T2D-Karlsson dataset is 100 amino acids, and the PD-Bedarf dataset is 30 amino acids. Thus we chose a fragment length of *l* = 30 for the experiments conducted in this paper.

Choice of the value of *k* need to be such that the chance of *k*-mers being shared by any two protein sequences in our gold standard datasets is minimized. In a study of 1,121 bacterial genomes (58) showed that for a *k*-mer length of *>* 20 nucleotides (*≥* 7 amino acids), over 96% of the nucleotide *k*-mers within an organism are unique and only less than 0.2% of the *k*-mers of length 25 nucleotides (*≥* 8 amino acids) are shared by any two organisms; the 25-mers have the same gene annotation in both genomes. Inspired by these results, we chose *k* = 8 for our experiments. With *k* = 8, we calculate the probability *p* of a random *k*-mer match within a dataset as follows (59):

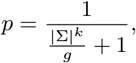

where, *|*Σ*|* is the alphabet size and *g* is the size of the dataset in total number of amino acids.

For EC-2192-DB (32,111,182 amino acids) this probability is 0.12%, and for KO-6282-DB (582,041,235 amino acids), this probability is 2.2%; both are sufficiently small. The effect of different k-mer and fragment lengths is shown in Supplementary Figure F2. For flexibility, *Carnelian* software takes fragment and k-mer lengths as input from the user.

### Performance evaluation metric

To evaluate the performance of *Carnelian*, we use the common F1-score metric – the harmonic mean of sensitivity (*ρ*) and precision (*π*). For each gold-standard functional label, *i*, we calculate *ρ* and *π* as follows:

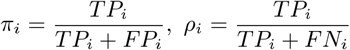

Here, *T P_i_* (True Positive) is the number of fragments binned correctly under label *i*; *FP_i_* (False Positive) is the number of fragments that do not have label *i* but are binned under label *i* by the classifier; and *FN_i_* (False Negative) is the number of fragments that belong to the bin of label *i* but were incorrectly assigned to some other bin. The overall F1-score of the entire binning problem can be computed by macro averaging, where F1-score for each bin, *F_i_*, is calculated first and then averaged over all bins as:

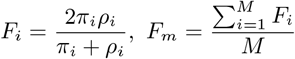

where *M* is the total number of unique functional labels. This measure has advantages over micro-averaged F1-scores because our measure gives equal weight to each label, regardless of how many examples of each the classifier has seen in the training set. Our measure is influenced by the classifier’s performance on relatively rare categories. The *Carnelian* software also reports micro-averaged precision, sensitivity, and F1-scores along with the macro-averaged ones.

### Benchmarking experiments

For the task of functional binning of simulated reads from proteins we used our gold standard dataset, EC-2192-DB for benchmarking experiments. We constructed the training set of *Carnelian* by drawing fragments of length, *l* = 30 from gold standard protein sequences such that every base has 5x coverage, and removed any duplicate fragments. Similarly, the test set was constructed by drawing 30 amino acid fragments from the gold standard proteins making sure that none of the fragments appeared in the training set.

For comparison with mi-faser, we constructed a DIAMOND database with protein sequences from our gold standard dataset. For Kallisto, we constructed an index from the gold standard dataset. Since a pre-compiled binary version of Kallisto expects nucleotide sequences as input, we back-translated the gold standard proteins using Backtranseq tool from the EMBOSS software suite (60) before indexing them. The same test set was used for all three tools. Kallisto was run with its “pseudobam” option to generate a bam file with the pseudo-alignment of fragments. Both mi-faser and Kallisto can place a fragment in possibly multiple bins; therefore, while looking for correctly annotated fragments, we consider a fragment as a true positive if atleast one of the labels predicted by the respective method matched ground truth. We reported macro-averaged F1-scores for all three methods. We ran *Carnelian* in both “default” and “precise” mode. In “default” mode, all the fragments are binned under the best possible label, whereas in “precise” mode the fragments are assigned to a bin only if the probability of the fragment belonging to that bin is greater a than user-specified probability threshold. All three methods were run on a single core of an Intel Xeon E5-2695 v2 x86 64 CPU @ 2.40 GHz. Note that, mi-faser and Kallisto have multithreading capabilities and runs very fast on parallel settings. See “Supplementary Text” for the exact commands used in running mi-faser and Kallisto.

For the task of binning novel proteins with known EC labels, we used EC-499-TR for training *Carnelian* as well as generating a DIAMOND database for mi-faser and index file for Kallisto. All three tools were tested on novel protein fragments from EC-499-TS as described above.

### Abundance Estimation

Since multiple reads can map to a single gene as well as multiple genes to a single read, depending on the lengths of the gene and the read, it is important to normalize the binned read/fragment counts (resp.) using both nominal read/fragment length (resp.) and average gene lengths per EC bin. Also, counts need to be normalized against the library size. Thus, we pursue the intuition behind transcripts per million (TPM) counts (34) and calculate effective read counts per bin similarly to that of TPM. Our effective counts matrix can be used in combination with any differential abundance estimation pipeline which can handle over-dispersed count data. Here, we perform differential abundance analysis using R packages – limma (61), voom (62), and edgeR (63). A Benjamini-Hochberg (BH) false discovery rate (FDR) corrected *p*-value less than 0.1 was used for testing significance. Additional fold-change thresholds were selected appropriate for each dataset. We mapped the significant ECs to KEGG pathways using the KEGG mapper tool (64) and determined significance of overlap using hypergeometric test (65).

### Functional analysis of disease metagenomes

For performing functional analysis on the T2D-Qin, T2D-Karlsson, and PD-Bedarf datasets, we applied *Carnelian* in “default” mode trained on EC-2192-DB with k-mer length *k* = 8 and fragment length *l* = 30. We also ran mi-faser on all the datasets with our gold standard ECs and compared the results. For the T2D-Qin dataset, we compared female T2D patients against female healthy controls and male T2D patients against matched male controls with both *Carnelian* and mifaser and compared our results. For the T2D-Karlsson dataset, after removing the samples that appear as outliers in either *Carnelian*- or mi-faser-derived functional profiles, we compared the following pairs of datasets (see Datasets) for biological relevance: IGT vs. NGT; T2D vs. NGT; and IGT vs. T2D. For the PD-Bedarf dataset, we compared the PD males with age-matched healthy males. Finally, for each dataset, we used the ECs that were significantly enriched/depleted to classify patients versus controls using a Random Forest (RF) classifier in a *k*-fold cross-validation setting (T2D: *k*=10, PD: *k*=5) to test the discriminatory power of the markers found by each method.

## Acknowledgements

We thank Yun William Yu and Hyunghoon Cho for useful discussions and comments.

## Author contributions

S.N. and B.B conceived the idea and designed the experiments. S.N. implemented *Carnelian* pipeline and performed the analysis. S.N. and B.B. wrote the manuscript.

